# CytokineFindeR: an R-package for benchmarking methods and databases for identifying cytokines

**DOI:** 10.1101/2025.09.30.679635

**Authors:** Jeffrey S. Tang, Asees Singh, Amrit Singh

## Abstract

Cytokines play a central role in disease but are hard to study due to their short half-life and low abundance. Computational approaches have utilized gene-expression for pinpointing cytokine drivers of diseases. We benchmark various cytokine identification methods and found little congruency between gene sets for the same ligand, and variable performance in identifying cytokines based on curated ligand-receptor interactions. CytoSig, a model-based approach, was generally better at identifying the correct disease cytokine but failed for certain cytokines such as IL-13. We developed CytokinefindeR, an R package that enables comparative analysis of cytokine detection across 10 databases and four computational approaches.

## Background

Cytokines are small messenger proteins that play important roles in coordinating the immune response in health and disease [1]. Cytokines include chemokines, interleukins, interferons and growth factors and have different roles in immune cell activation, differentiation and proliferation [1]. Cytokines have pleiotropic effects that can be dependent on the cellular environment making it difficult to study them in disease conditions [1]. Further, quantification of cytokines is difficult due to their low abundance, transient expression, and short half-life. Cytokines can be measured using Enzyme-Linked Immunosorbent Assays (ELISAs) or multiplex cytokine assays such as bead-based or antibody-based arrays. However, as no gold standard exists for measuring cytokines, considerable assay-to-assay variation has been observed in the literature. For example, cytokine data from 4,000 patients hospitalized with COVID-19 indicated significant differences in quantification between Clinical Laboratory Improvement Amendments (CLIA)-accredited vendors [2]. In contrast to directly measuring cytokine activity, cellular gene-expression can be used to identify the driver cytokines for a given disease, either using ligand-receptor interactions (LRI) [3] or using a signature-matrix that captures cytokine-specific gene-expression profiles [4]. LRIs can be used in gene set enrichment analysis [5] where the ligand is the “pathway” and its receptors are the “pathway gene-members” in order to identify enriched cytokines. Gene summarization methods[6] can also be used with LRIs to compute ligand activity scores per sample, after which differential expression analysis may be performed to identify ligands whose levels significantly differ between conditions. Recently a predictive model called Cytokine Signaling Analyzer (CytoSig), was developed using cytokine-specific perturbation transcriptomics data to predict the sample-level activity of 43 cytokines [4]. To our knowledge, no previous benchmarking has been performed to compare the various computational approaches for cytokine identification. We leverage many LRI databases and a range of cytokine perturbation and anti-cytokine treatment transcriptomics datasets in order to compare LRI-based methods and CytoSig (a predictive model that does not use LRIs). We provide all these approaches for users to apply to their own datasets through an R-package called CytokineFindeR.

## Results and discussion

Given a gene-expression dataset (*N samples x P genes*) and vector of labels for the *N samples* (e.g. control/disease), CytokineFindeR can be used to determine statistical significance of cytokines based on LRI-based methods as well as a reimplementation of the ridge regression model of CytoSig (Fig. A). CytokineFindeR contains 10 curated LRI databases: BaderLab, LIANA+, OmniPath, NicheNet, scDiffCom, CellChat, iTalk, FANTOM5, CytokineLink and combined (see Table S1 for details). We used CytokineFindeR to benchmark 30 LRI-based approaches (10 databases x 3 LRI-based methods; gene set enrichment analysis and two summarization techniques, principal component analysis and gene set variation analysis) and CytoSig (either using the online tool or our reimplementation in R) using both cytokine perturbation and treatment studies (Fig. 1B). Cytokine perturbation studies included neutrophils stimulated with IFNG, IL-4, IL-13 and endothelial cells stimulated with TNF, whereas anti-cytokine treatment studies related to the inhibition of TNF, IL-1B, IL-12B, IL-13 and IL-17A across various disease conditions (Table S2).

**Figure 1.**
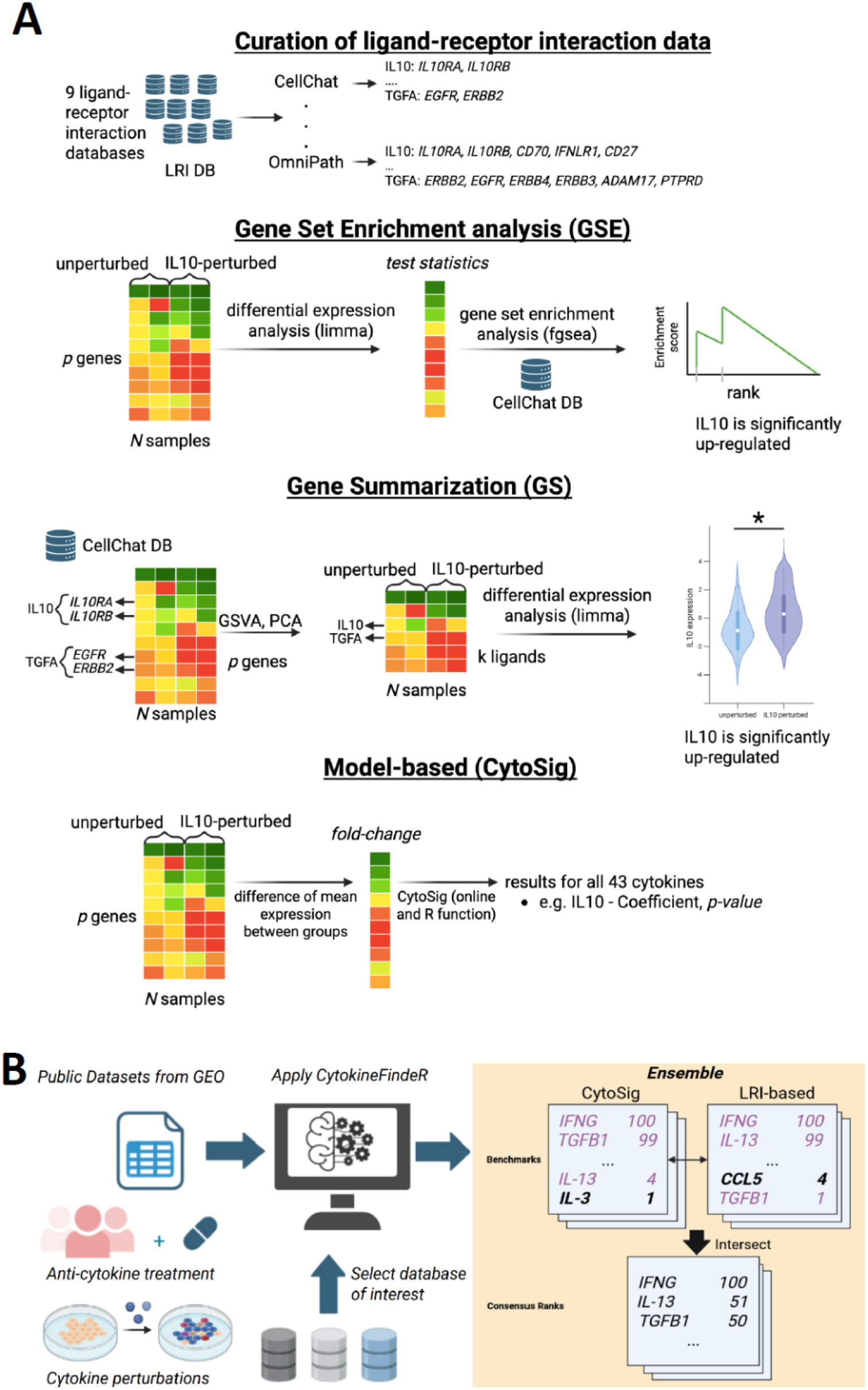
Databases and methods implemented in the CytokineFindeR R-package and its benchmarking using cytokine perturbation anti-cytokine treatment gene-expression data. A. ligand-receptor database curation and methods for cytokine identification. Ligand-receptor interaction (LRI) pairs were curated from 9 sources. Methods were categorized into three classes: gene set enrichment (GSE), gene summarization (GS), and model-based (CytoSig). Both GSE and GS methods utilize LRI pairs and are coupled with differential expression analyses in order to identify differentially expressed ligands. CytoSig uses log_2_ fold-change as an input and computes the statistical significance for each cytokine. B. CytokineFindeR was used to benchmark cytokine identification methods by ranking methods based on their ability to rank the “target” cytokine of each study relative to the 43 tested cytokines (since CytoSig is restricted to 43 cytokines). An ensemble approach based on the significance of LRI-based methods and CytoSig is also included. Created in BioRender.com

Fig. 2A depicts the number of ligands in each ligand-receptor interaction (LRI) database, with CytokineLink consisting of the least number of ligands (64 including 43 high-confidence cytokines) and BaderLab consisting of the greatest number of ligands (2293). Given the large range of ligands contained across the 9 databases, we sought to determine the overlap coefficient (OLC, ie. how much a smaller set is contained within a larger one) between the databases. The OLC with respect to the ligands in each database ranged between 0.48 to 1.00. CellChat was the most distinct from other databases (OLC ranged between 0.58-0.90) whereas CytokineLink had most ligands in common with the other databases (OLC ranged between 0.90-0.99) with the exception of CytoSig (Fig. 2B). The Jaccard Similarity Score (JSS) was calculated on a per-ligand basis to assess the similarity between receptors of the same ligand across databases (see Methods for details). We found 29 common ligands across all databases and assessed receptor annotation similarity. On average, the receptors for the 29 ligands were dissimilar except for FANTOM5 and iTALK which had a JSS of 1. Amongst the LRI databases, BaderLab consisted of receptor gene symbols that overlapped the least with all other databases. Upon closer inspection these “receptors” consisted of other molecules such as the ligand itself (e.g. TNF was one of the receptor genes listed for itself). The “receptor” genes determined for CytoSig had very little overlap across all LRI databases (Fig. 2C).

**Figure 2.**
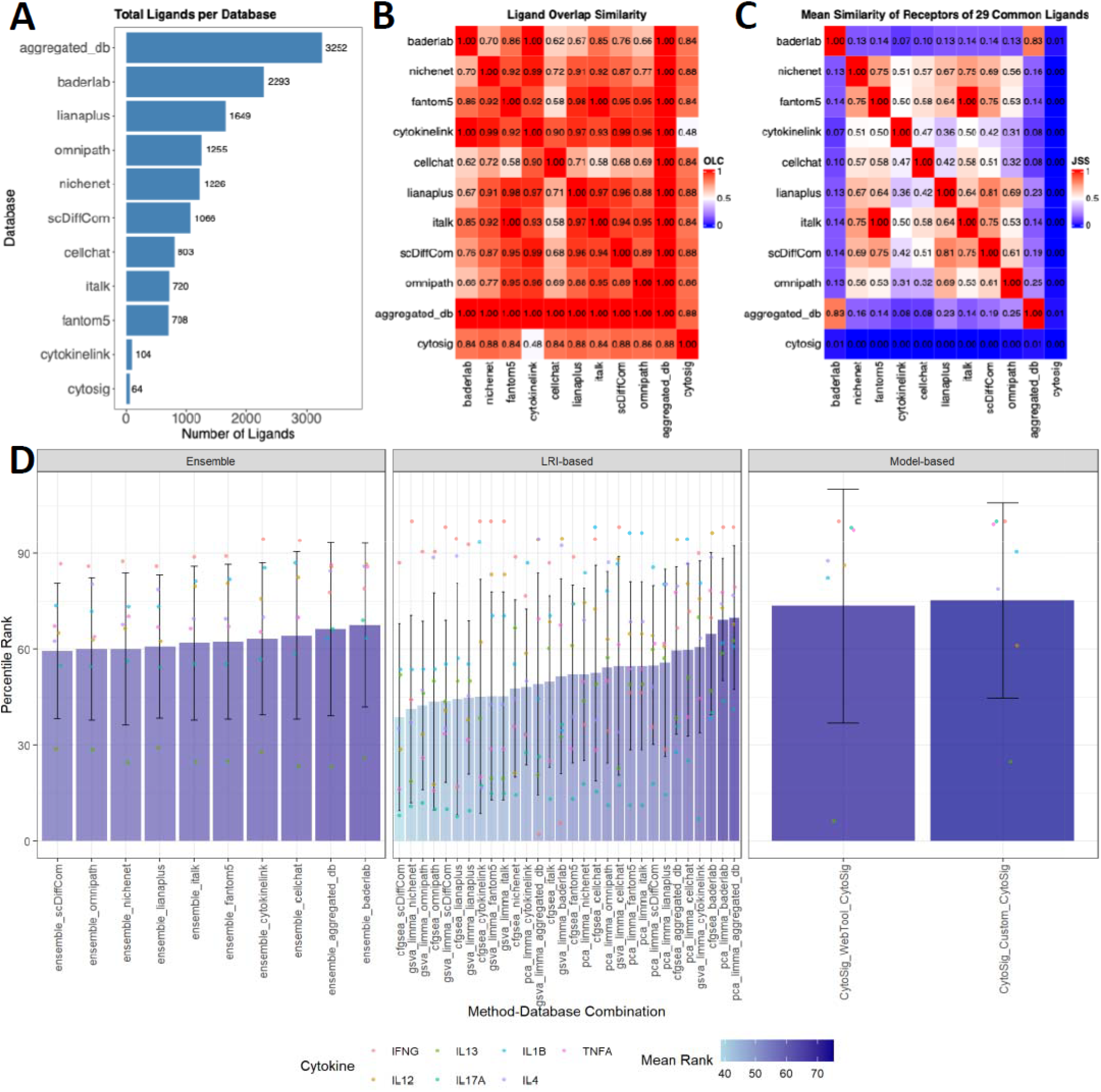
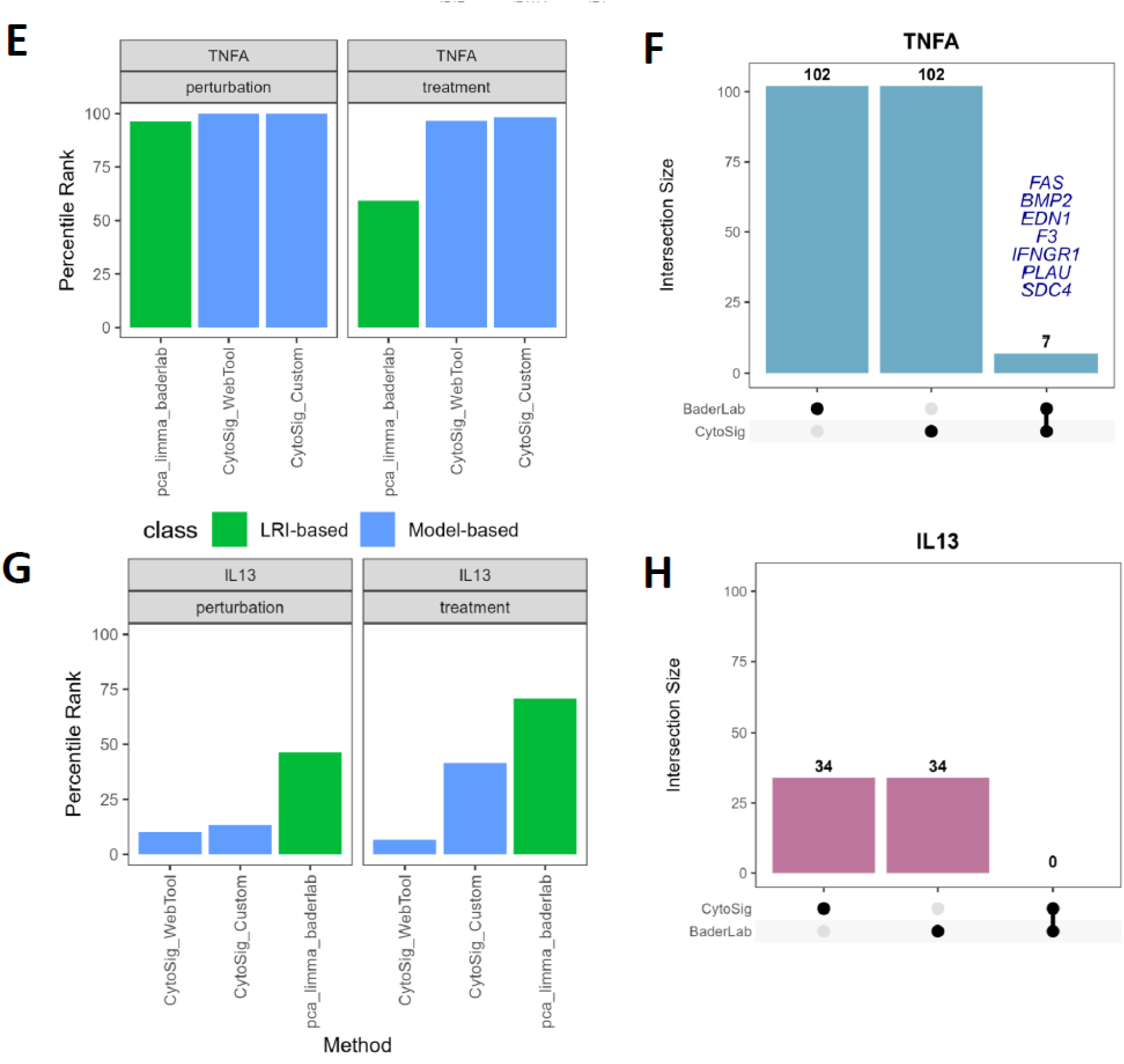
Benchmarking LRI-based and model-based methods using cytokine perturbation and anti-cytokine treatment gene-expression data. A. Number of ligands in each LRI database. B. Overlap coefficients of ligands recorded across 9 databases, all databases combined (aggregated_db) and CytoSig cytokines. C. Mean Jaccard Similarity Score (JSS) of 28 common ligands found across the databases. D. Benchmark observations of targeted-cytokine perturbation and treatment experiments: The mean ± standard deviation ranking across the various LRI-based, method-based and ensemble-based methods. E. Rank of TNF based on the top-performing LRI-method-database combination (pca_limma_baderlab) and CytoSig for both perturbation and treatment datasets. The model-based approach, CytoSig outperforms the LRI-based method. F. 7 receptor-genes overlapped between the BaderLab database and CytoSig signature matrix for TNF. G. Rank of IL-13 based on the top-performing LRI-method-database combination (pca_limma_baderlab) and CytoSig for both perturbation and treatment datasets. The LRI-based method out-performs the model-based approach, CytoSig. H. No receptor-genes overlapped between BaderLab database and CytoSig signature matrix for IL-13.

Benchmarking of the LRI methods was restricted to the ligands that overlapped with CytoSig (see Table S3 for the number of cytokines considered for each database). Fig. 2D depicts the ability of each method to correctly identify the true cytokines from the 29 tested cytokines across the four stimulation and 5 treatment datasets. CytoSig had an average ranking of 73.5±36.5 and 75.3±30.5 for the web-tool and R-implementation respectively. The top-performing LRI method was pca_limma with an average rank of 69.2±19.2 and 69.8±22.4 when used with BaderLab and all databases combined respectively. Cytokines present in both the simulation and treatment datasets indicated superior performance of CytoSig over pca_limma_baderlab for TNF, whereas pca_limma_baderlab outperformed CytoSig for IL-13 (Fig. 2E, 2G). Interestingly 7 genes contributing strongly to CytoSig’s prediction of TNF overlapped with prior known “receptors” of TNF as part of the BaderLab database (Fig. 2F), where no overlap was identified for IL-13 (Fig. 2H).

Our results indicate that the information contained within existing LRI databases is variable and can significantly impact downstream analyses. Principal component analysis is superior in imputing ligand expression using gene-expression profiles of receptor genes, compared with other summarization methods such as GSVA. Although CytSig outperformed existing LRI-based method for many cytokines, it predictively ability is dependent on the amount of training data used to develop the model. For example, many studies specific to TNF whereas much less studies on IL-13 were included when training the original CytoSig model [4]. This may suggest why performance of CytoSig in other evaluated datasets is much better than IL-13 when compared to LRI-based. The authors of CytoSig mentioned ascertainment biases and provide a breakdown of study counts which reported that IL-13 was trained using under 50 studies [4]. This highlights how training data limitations can affect method performance on specific cytokines. Similarly, the LRI databases underlying our benchmarked methods were each curated from different experimental datasets and annotation approaches, which may contribute to the performance variability we observe across methods and cytokines. Our benchmarking results suggest that using PCA on LRI data from the BaderLab may be a useful approach for identifying cytokines in cases where no predictive model exists for estimate cytokine levels or if little data was present for a particular cytokine during training.

Despite advances in omics technologies, barriers such as software complexity and usability continue to limit accessibility and standardization of procedures. This underscores the need for open, dynamic benchmarking frameworks—such as CytokineFindeR—that facilitate continuous, transparent evaluation of tool performance. Beyond cytokine detection, CytokineFindeR’s modular design allows users flexibility in method and LRI database selection.

## Methods

### Study Selection of Publicly Available Datasets

Publicly available datasets were curated from the Gene Expression Omnibus (GEO) as shown in Fig. 1B. Dataset curation was conducted for 2 human-centric experimental designs with either normal cells or a range of immune-related conditions to be assessed individually with preprocessed gene expression or counts matrices for our benchmarking framework. We narrowed search space to cytokine-related differential cell states based on either perturbation or targeted therapeutic experimental designs. Post-dataset curation, studies were filtered against class imbalance between differential (normal vs. perturbed or treatment time points vs baseline) conditions and smaller sample sizes (n < 2). We cross-referenced published training data from the selected data-driven model-based approach to ensure no overlap in selected benchmarking datasets for validation purposes. Nine studies were selected for the current benchmarking report. Variation in study design such as condition, timepoints, and paired sample consideration should be handled during pre-processing and must be specified in the workflow. Table 1 provides a comprehensive overview of all studies selected. Given that all included studies assessed gene expression based on cytokine targets, this provides us with a degree of known ground truth.

#### Cell-Isolate Perturbation Experiments

Chemical perturbation studies pertaining to different types of cytokine exposures against cell-isolates to illicit a response are useful to gain insight on pathways in an isolated environment. Several perturbational studies assessed changes across different time points and conditions using peripheral blood mononuclear cells (PBMCs) in response to target cytokines. We referenced a human neutrophil study isolated from whole blood that perturbed activity of IL-13, IL-4, and IFNG between 2 time points and a separate study that stimulated TNF on human umbilical vein endothelial cells between 2 time points.

#### Targeted Therapy Studies

We collected 5 targeted treatment studies following similar selection criteria. Treatment-based studies looked at specific immune disease conditions and their response to targeted drugs relevant to cytokines. By applying benchmarking on treatment studies, we can also assess cytokine activity detection across a variety of disease conditions compared to perturbation studies which measure activity of normal cells.

### Cytokine identification methods

We report on transcriptomics methods coupled with ligand-receptor interactions (LRIs) to provide insight into cytokine activity. We categorized methods into two approaches: “data-driven model-based” and LRI-based methods. Data-driven model-based approaches require explicit training on large external datasets before application to new data for molecular activity inference. In contrast, LRI-based methods use pre-defined gene sets such as with gene set enrichment (GSE) or gene summarization (GS) approaches, applied directly post-differential expression analyses without prior model training. We also include principal component analysis (PCA) as a GS method. Methods selection was determined by previous benchmark findings, popularity, and ease of integration [Xie *et al*., 2021], focusing on two well-established tools widely used for pathway activity inference. Our approach leverages LRI databases to establish receptor gene sets that serve as proxies for cytokine activity (see Fig. 1A for differences in methods implemented). This suite of conventional R tools forms a baseline for comparison, with flexibility for users to customize and integrate additional exploratory methods.

#### Gene Set Enrichment

Gene set enrichment analysis (GSEA) has become a household tool for inferring enriched pathways, ranking their significance by association to pre-defined gene sets provided to the tool [Subramanian *et al*, 2005]. Similarly, we can provide sets of ligands or cytokines of interest explained by their receptor genes based on a reference source to explain their activity. We chose a popular and faster iteration of GSEA (fgsea) which allows for decreased sampling requirements whilst maintaining the integrity of accurate p-values of the original implementation [Korotkevich *et al*, 2019].

#### Gene Summarization

Gene summarization is an approach that reduces complexity of gene expression dimensionality. These approaches involve reducing high-dimensionality of gene expression to a smaller set of key features such as using PCA or transforming gene-level data into gene lists that represent specific signatures such as pathways using Gene Set Variation Analysis (GSVA). The distinction from GSE is that it does not require pre-ranking of gene lists which are commonly generated through differential expression analysis first [Hanzelmann *et al*, 2013]. For our use case, based on a transformation of gene expression representation across the sets of receptor genes first, we can determine significantly enriched ligands.

#### Data-driven Model-Based Approach

Among current computational tools dedicated to cytokines, Cytokine Signaling Analyzer (CytoSig) uniquely offers a systematic approach for summarizing cytokine signatures based on a corpus of cytokine-driven studies. A web tool is readily available for use which requires differentially expressed log2 fold changes between two conditions as input to generate a signature of cytokines. CytoSig is a ridge regression model that utilizes a gene scoring matrix to account for co-expression of genes trained on 20,591 transcriptome profiles from 962 experiments [Jiang *et al*, 2021]. The β coefficients which capture co-expression were published in their source code and enabled us to implement an R version of the tool for integration into CytokineFindeR. We also reported on benchmarking results using the web tool for transparency (https://cytosig.ccr.cancer.gov/). CytoSig was the gold standard method to compare against due to its reliability in characterizing common cytokines occurring in the literature and volume of training data used.

### Database Selection and Comparison

We aggregated 9 reference databases for benchmarking LRI-based cytokine activity predictions across an assortment of recently published and actively maintained databases. LRI annotations were compiled across selected databases. Some databases were derived from scRNAseq data such as LIANA+ [Dimitrov *et al*, 2024], CellChatDBv2 [Jin *et al*, 2024], scDiffCom [Lagger *et al*, 2023], and iTALK [Wang *et al*, 2019]. We collapsed cell-level annotations, pooling all receptor associations to the ligand as we did not include the scope of cell-level resolutions. To gain insight on database variation, we performed a meta-analysis of concordance using overlap coefficients (OLC), defined as the size of the intersection divided by the smaller size of the two database sets, to assess and quantify ligand overlap. The intersect of common ligands (n = 28) across all databases was identified to assess receptor across all databases and we performed Jaccard Similarity (JS), defined as the size of the intersection divided by the union of two database sets, as the metric to evaluate similarity. We used OLC scores for ligand comparisons due to the vastly different ligand set sizes for each database, which could range from 104 to 3252 ligands; thus limiting the comparison relative to the smallest set of ligands yields better comparisons. Furthermore, we included CytoSig in the database comparison by selecting the top genes ranked by absolute β coefficients, limited to match the receptor set size of each reference database for a given ligand to calculate JS scores.

## Data and code availability

The gene-expression datasets used for benchmarking can be found on Zenodo. The GitHub code used to reproduce all analyses can be found here (https://github.com/CompBio-Lab/cytokinefinder_analysis) whereas the CytokineFinder R-package can be found here (https://github.com/CompBio-Lab/cytokineFinder).

## References

1. Mallick S, Duttaroy AK, Bose B. A Snapshot of Cytokine Dynamics: A Fine Balance Between Health and Disease. J of Cellular Biochemistry [Internet]. 2025 [cited 2025 July 22];126. Available from: https://onlinelibrary.wiley.com/doi/10.1002/jcb.30680

2. Wang SY, Takahashi T, Pine AB, Damsky WE, Simonov M, Zhang Y, et al. Challenges in interpreting cytokine data in COVID-19 affect patient care and management. PLoS Biol. 2021;19:e3001373.

3. Türei D, Korcsmáros T, Saez-Rodriguez J. OmniPath: guidelines and gateway for literature-curated signaling pathway resources. Nat Methods. 2016;13:966–7.

4. Jiang P, Zhang Y, Ru B, Yang Y, Vu T, Paul R, et al. Systematic investigation of cytokine signaling activity at the tissue and single-cell levels. Nat Methods. 2021;18:1181–91.

5. Korotkevich G, Sukhov V, Budin N, Shpak B, Artyomov MN, Sergushichev A. Fast gene set enrichment analysis [Internet]. Cold Spring Harbor Laboratory; 2016 [cited 2025 July 22]. Available from: http://biorxiv.org/lookup/doi/10.1101/060012

6. Badia-i-Mompel P, Vélez Santiago J, Braunger J, Geiss C, Dimitrov D, Müller-Dott S, et al. decoupleR: ensemble of computational methods to infer biological activities from omics data. Kuijjer ML, editor. Bioinformatics Advances [Internet]. 2022 [cited 2025 July 22];2. Available from: https://academic.oup.com/bioinformaticsadvances/article/doi/10.1093/bioadv/vbac016/6544613

