## Supplementary material for "CytokineFindeR: an R-package for benchmarking methods and databases for identifying cytokines": Table S1, Table S2, Table S3

**Supplementary Information**

**Table S1.** Ligand-interaction databases

| **Databases** | **Number of ligands** | **Total number of receptors (ligand-related genes)** | **Reference** |
| --- | --- | --- | --- |
| Baderlab | 2293 | 2492 | https://baderlab.org/CellCellInteractions?action=AttachFile&do=get&target=receptor_ligand_interactions_mitab_v1.0_April2017.txt.zip |
| NicheNet | 1226 | 1067 | [1] |
| FANTOM5 | 708 | 691 | https://fantom.gsc.riken.jp/5/ |
| CITEdb | 62 | 59 | [2] |
| CytokineLink | 104 | 101 | [3] |
| CellChat | 800 | 720 | [4]  https://github.com/jinworks/CellChat |
| LIANA+ | 1649 | 1410 | [5] |
| iTALK | 720 | 699 | [6] |
| scDiffCom | 1066 | 954 | [7] |
| OmniPath | 1255 | 1279 | [8] |
| AggregatedDB* | 3252 | 3387 |  |

*all databases combined

**Table S2.** Overview of all studies selected for benchmarking.

| Design | Cytokine | Data | Biospecimen | Description |
| --- | --- | --- | --- | --- |
| Perturbation | *IFNG* | GSE135425 | Blood-isolated neutrophils | Effect of *IFNg* on gene expression on human neutrophil isolates in 3 controls and 3 stimulated samples from healthy donors [9] |
|  | *IL-4* | GSE135421 | Blood-isolated neutrophils | Effect of *IL-4* on gene expression on human neutrophil isolates in 3 controls and 3 stimulated samples from healthy donors [9] |
|  | *IL-13* | GSE135426 | Blood-isolated neutrophils | Effect of *IL-13* on gene expression on human neutrophil isolates in 3 controls and 3 stimulated samples from healthy donors [9] |
|  | *TNF* | GSE179478 | Endothelial cell line | Human umbilical vein endothelial cells (HUVEC) stimulated with *TNF* for 24 hours compared to basal condition [10] |
| Treatment | *TNF* | GSE92415 | Colon mucosal biopsy | Ulcerative colitis patient GEPs treated with golimumab (anti-*TNF*) from baseline and week 6 [11] |
|  | *IL-1B* | GSE80060 | Whole blood | Canakinumab (human anti-*IL-1B*) treatment effects on whole blood of juvenile idiopathic arthritis patients [12] |
|  | *IL-12B* | GSE106992 | Skin biopsy | Ustekinumab profiling for *IL-12* and *IL-13* changes at baseline and week 12 [13] |
|  | *IL-13* | GSE277961 | Skin biopsy | Inhibition of IL-13 in atopic dermatitis with the use of tralokinumab from baseline and week 2 [14] |
|  | *IL-17A* | GSE226244 | Whole tissue of psoriasis skin lesions | *IL-17a* blockade treatment on psoriasis patients measured at 12 and 24 hours [15] |

Table S3: Number of cytokines used for benchmarking each database

| Database | Overlap with CytoSig cytokines |
| --- | --- |
| BaderLab | 53-55 |
| NicheNet | 52-56 |
| FANTOM5 | 54 |
| CITEdb* | 10 |
| CytokineLink | 29-31 |
| CellChat | 51-53 |
| LIANA+ | 53-56 |
| iTALK | 54 |
| scDiffCom | 49-56 |
| OmniPath | 51-55 |

* CITEdb was removed from all further analyses as there were too little overlapping cytokines to provide meaningful performance ranking.

**References**

1. Browaeys R, Saelens W, Saeys Y. NicheNet: modeling intercellular communication by linking ligands to target genes. Nat Methods. 2020;17:159–62.

2. Shan N, Lu Y, Guo H, Li D, Jiang J, Yan L, et al. CITEdb: a manually curated database of cell–cell interactions in human. Bioinformatics. 2022;38:5144.

3. Olbei M, Thomas JP, Hautefort I, Treveil A, Bohar B, Madgwick M, et al. CytokineLink: A Cytokine Communication Map to Analyse Immune Responses—Case Studies in Inflammatory Bowel Disease and COVID-19. Cells. 2021;10:2242.

4. Jin S, Plikus MV, Nie Q. CellChat for systematic analysis of cell–cell communication from single-cell transcriptomics. Nat Protoc. 2025;20:180–219.

5. Dimitrov D, Schäfer PSL, Farr E, Rodriguez-Mier P, Lobentanzer S, Badia-i-Mompel P, et al. LIANA+ provides an all-in-one framework for cell–cell communication inference. Nat Cell Biol. 2024;26:1613–22.

6. Wang Y, Wang R, Zhang S, Song S, Jiang C, Han G, et al. iTALK: an R Package to Characterize and Illustrate Intercellular Communication [Internet]. bioRxiv; 2019 [cited 2025 Jul 27]. p. 507871. Available from: https://www.biorxiv.org/content/10.1101/507871v1

7. Lagger C, Ursu E, Equey A, Avelar RA, Pisco AO, Tacutu R, et al. scDiffCom: a tool for differential analysis of cell–cell interactions provides a mouse atlas of aging changes in intercellular communication. Nat Aging. 2023;3:1446–61.

8. Türei D, Korcsmáros T, Saez-Rodriguez J. OmniPath: guidelines and gateway for literature-curated signaling pathway resources. Nat Methods. 2016;13:966–7.

9. L K, T S, Z O, R S, Ha Y, Is J. IL-4, IL-13 and IFN-γ -induced genes in highly purified human neutrophils. Cytokine [Internet]. 2023 [cited 2025 Jul 27];164. Available from: https://pubmed.ncbi.nlm.nih.gov/36809715/

10. Michaeli JC, Albers S, de la Torre C, Schreiner Y, Faust S, Michaeli T, et al. Gene regulation for inflammation and inflammation resolution differs between umbilical arterial and venous endothelial cells. Sci Rep. 2023;13:16159.

11. Wj S, Bg F, C M, H Z, R S, J J, et al. Subcutaneous golimumab induces clinical response and remission in patients with moderate-to-severe ulcerative colitis. Gastroenterology [Internet]. 2014 [cited 2025 Jul 27];146. Available from: https://pubmed.ncbi.nlm.nih.gov/23735746/

12. Ah B, Aa G, N W, Hi B, P Q, R B, et al. Early changes in gene expression and inflammatory proteins in systemic juvenile idiopathic arthritis patients on canakinumab therapy. Arthritis research & therapy [Internet]. 2017 [cited 2025 Jul 27];19. Available from: https://pubmed.ncbi.nlm.nih.gov/28115015/

13. C B, K L, S G, K H, A C, I N, et al. Modulation of inflammatory gene transcripts in psoriasis vulgaris: Differences between ustekinumab and etanercept. The Journal of allergy and clinical immunology [Internet]. 2019 [cited 2025 Jul 27];143. Available from: https://pubmed.ncbi.nlm.nih.gov/30703387/

14. Tandon R, Harder I, Stölzl D, Hübenthal M, Sander N, Hartmann J, et al. Tralokinumab Treatment of Atopic Dermatitis Induces a Progressive Transcriptomic Response. J Invest Dermatol. 2025;145:1643-1652.e13.

15. J K, J L, X L, N K, D R, I C, et al. Multi-omics segregate different transcriptomic impacts of anti-IL-17A blockade on type 17 T-cells and regulatory immune cells in psoriasis skin. Frontiers in immunology [Internet]. 2023 [cited 2025 Jul 27];14. Available from: https://pubmed.ncbi.nlm.nih.gov/37781383/
